# Unravelling the microplastic menace: different polymers work in synergy to increase bee vulnerability

**DOI:** 10.1101/2024.02.08.579429

**Authors:** Federico Ferrante, Elisa Pasquini, Federico Cappa, Lorenzo Bellocchio, David Baracchi

## Abstract

Microplastics (MPs) are growing and ubiquitous environmental pollutants and represent one of the greatest contemporary challenges caused by human activities. Current research has predominantly examined the singular toxicological effects of individual polymers, neglecting the prevailing reality of organisms confronted with complex contaminant mixtures and potential synergistic effects. To fill this research gap, we investigated the lethal and sublethal effects of two common MPs, polystyrene (PS - 4.8-5.8 μm) and poly(methyl methacrylate) (PMMA - 1-40 μm), and their combination (MIX), on the pollinating insect *Apis mellifera*. For each treatment, we evaluated the oral toxicity of two ecologically relevant and one higher concentration (0.5, 5 and 50 mg/L) and analysed their effects on the immune system and worker survival. As immune activation can alter the cuticular hydrocarbon profile of honey bees, we used gas chromatography-mass spectrometry (GC-MS) to investigate whether MPs lead to changes in the chemical profile of foragers and behavioural assay to test whether such changes affect behavioural patterns of social recognition, undermining overall colony integrity. The results indicate a synergistic negative effect of PS and PMMA on bee survival and immune response, even at ecologically relevant concentrations. Furthermore, alterations in cuticle profiles were observed with both MPs at the highest and intermediate concentrations, with PMMA being mainly responsible. Both MPs exposure resulted in a reduction in the abundance of several cuticular compounds. Hive entry guards did not show increased inspection or aggressive behaviour towards exposed foragers, allowing them to enter the colony without being treated differently from uncontaminated foragers. These findings raise concerns not only for the health of individual bees, but also for the entire colony, which could be at risk if contaminated nestmates enter the colony undetected, allowing MPs to spread throughout the hive.

## Introduction

Over the last 70 years, the world has become increasingly dependent on plastics (Hale et al., 2020). Their versatility, stability, light weight, and low production costs made them the most ubiquitous materials in contemporary history (Cole et al., 2011). As a result of this sustained and massive use, the natural environment is heavily contaminated with plastic waste and debris (Zheng et al., 2023), including microplastics (MPs), defined as polymer particles smaller than five mm in diameter (Frias & Nash, 2019; C. Wang et al., 2021). Due to their strong hydrophobicity, small size, large specific surface area and stable chemical properties, these pollutants can accumulate, migrate, and diffuse in a wide range of environmental matrices, including air, soil, and water (Ng et al., 2018; Priya et al., 2022; Rochman, 2018; Yu et al., 2022).

The exponential contamination by MPs is not only a growing concern for environmental pollution but also a threat to the health of living organisms. These minute particles can be ingested by various species through the food chain and cause a range of detrimental effects, including reduced growth, oxidative stress, intestinal damage, cardiotoxicity, immunotoxicity and reproductive toxicity (Chang et al., 2022; Zheng et al., 2023; Zhou et al. 2023). To date, most studies have focused on marine environments (Andrady, 2011; Botterell et al., 2019; Eerkes-Medrano et al., 2015; Li et al., 2020; de Souza Machado et al., 2018), but the attention has recently expanded to terrestrial ecosystems and the associated risks to land-dwelling organisms (Zeb et al., 2024). It has been estimated that MPs terrestrial contamination could be far greater than in oceans (Horton et al., 2017). Furthermore, soil contamination is considered potentially more dangerous than aquatic contamination due to its direct effects on food chains, wild plants and animals, crops and livestock (Dissanayake et al., 2022; Wang et al., 2021). Strikingly, MPs can affect terrestrial organisms that mediate essential ecosystem services and functions, such as pollinators (de Souza Machado et al., 2018).

The western honey bee, *Apis mellifera*, serves as an exemplary model organism for terrestrial ecotoxicology, due to its critical role as a primary pollinator for both wild and cultivated plants. (Gallai et al., 2009; Moritz et al., 2010). As bees forage, they are constantly exposed to environmental pollutants, including MPs, as demonstrated by recent studies that found MPs in honey (Diaz-Basantes et al., 2020; Liebezeit & Liebezeit, 2013) and in the bee cuticle (Deng et al., 2021; Edo et al., 2021). Adding to their vulnerability, bees possess a limited number of genes encoding detoxification enzymes (Claudianos et al., 2006). For these reasons, the honey bee is considered an excellent bioindicator for monitoring trace contaminations in the environment (Badiou-Bénéteau et al., 2013; Wang et al., 2021). Moreover, honey bees are eusocial insects, which offers the unique advantage for studying the effects of pollutants at different levels of biological organization. Despite the advantages, only a limited number of studies have explored the potential hazards associated with MPs exposure in bees. Existing research indicates reduced bees’ feeding rate (Al Naggar et al., 2023), modified expression of antioxidative, detoxification, and immune system-related genes in young bees (Wang et al., 2021), impacted learning and memory in foragers (Balzani et al., 2022; Pasquini et al. 2023), and increased the susceptibility to viral infections (Deng et al., 2021). The comprehensive evidence outlined above highlights the multifaceted effects of MPs on honey bees, signalling a potential threat to both individual bees and the colony as a whole. In recognition of this, ecotoxicology must adopt a holistic approach, assessing not only the effects of these environmental contaminants on the survival and physiology of individual bees, but also their potential influence on colony interactions. This broader perspective is consistent with recent research on other species of social insects and contaminants (Cappa et al., 2019, 2024; Gill et al., 2012; Michelangeli et al., 2022). Moreover, despite the growing interest towards MPs pollution, current studies still present further limitations, focusing solely on the ecotoxicological effects of single polymers. It is widely recognised that in the environment organisms are often exposed to combinations of contaminants, which could likely result in additive or synergistic effects (Siviter et al., 2021). Due to the significant gap in the risk assessment of these mixtures, the real impact of MPs remains inadequately addressed and their actual hazard is most probably underestimated. To start bridging these knowledge gaps, in the present study, we adopted a more holistic approach in ecotoxicology by assessing the impact of two environmentally prevalent MPs, polystyrene (PS) and polymethylmethacrylate or plexiglass (PMMA), both singly and in combination, on both individual and social traits in honey bees. In particular, we assessed the effects of exposure to MPs on the survival, immune system and chemical profile of individual bees, and the impact of such potential alterations on the colony’s ability to detect contaminated foragers and prevent colony exposure by collectively rejecting them.

We chose to study these aspects because the individual health status of an individual bee is often mirrored in its cuticular hydrocarbons (CHCs) profile. CHCs play a crucial role in recognition processes at the colony level in honey bees and other social insects (Bortolotti & Costa, 2014; Van Zweden & d’Ettorre, 2010). In particular, activation of the immune system can induce modifications in the CHCs of individuals (Baracchi et al., 2012; Richard et al., 2008, 2012; Cappa et al., 2016). These modifications in CHCs can be used by colony members to detect, quarantine, or remove diseased or parasitised individuals within the nest (Baracchi et al, 2012; Cappa et al., 2016, 2019; Clancy, 1996, Milutinović & Schmitt, 2022). Therefore, if exposure to MPs affects individual health and survival, and these adverse effects are reflected in the chemical profile, we expect contaminated bees to be rapidly identified and rejected by guard bees at the hive entrance. This proactive response aims to prevent colony exposure and protect colony integrity. Our ultimate goal is, therefore, to provide reliable insights into how synergistic exposure to MP pollutants can affect free-living organisms at multiple levels, undermining their survival and ecosystem services.

## Material & Methods

### MPs Solutions

Experimental groups were exposed to polystyrene (PS), Polymethyl-methacrylate/plexiglass (PMMA) or a combination of both (hereafter “MIX”) to assess any potential additive or synergistic effects. MPs solutions were prepared using PS (PSMS-1.07, 4.8-5.8 μm, Cospheric LLC, USA) and PMMA (PMPMS-1.2, 1-40 μm, Cospheric LLC, USA) microspheres at three different concentrations (0.5, 5 and 50 mg/L) in purified water with 50% (w/w) sucrose. These concentrations were designated as low (L), medium (M) and high (H). L and M solutions represented the realistic range of variability found in the natural environment (Chae et al., 2018; Eltemsah and Bohn, 2019; Wang et al., 2021), while H a higher concentration to assess acute intoxication effects. The 50% (w/w) sucrose solution used for the control bees did not contain MPs.

### Survival Assay

We first investigated the effects of chronic oral exposure to PS, PMMA, and the MIX on the survival and daily sucrose intake of confined honey bees collected from an experimental apiary at the Department of Biology, University of Florence (Italy). Following established toxicological research protocols for *A. mellifera* (Williams et al., 2013), bees were collected and housed, in groups of 10 individuals, in 120 ml Plexiglass cages. Each cage was then equipped with a 20 ml syringe containing either PS, PMMA, MIX sucrose solution, or solely a plain sucrose solution. The syringes were modified by removing the luer cone to allow easy access to the food while preventing any spillage of the sugar solution (Carlesso et al., 2020). We assigned 40 cages to each group (PS, PMMA, MIX), of which 10 served as controls, 10 received the high concentration (H), 10 the medium concentration (M) and 10 the low concentration (L). Overall, we tested a total of 120 bees cages. To account for potential solution evaporation, an additional syringe was positioned in an empty cage to mimic the experimental conditions throughout the study. Daily survival rates were recorded by counting the number of dead bees, and the amount of sucrose consumed was measured by weighing the syringes each day. Daily sucrose intake was adjusted based on the number of live bees in each cage on any given day (Carlesso et al., 2020). The cages were maintained in darkness at room temperature (23 ± 2 °C) and humidity levels of (50 ± 10%) throughout the experiment.

### Immune Challenge Assay

Forager bees were collected from six hives of the same experimental apiary. We compared the ability of foragers exposed to different concentrations of PS, PMMA, and MIX to clear bacterial cells from their haemolymph (i.e., bacterial clearance) by injecting bees with the Gram-negative bacteria *Escherichia coli*, an immune elicitor used to test immunocompetence in insects (Yang and Cox-Foster 2005; Cappa et al. 2020, 2022). We measured bacterial clearance as a good proxy of workers immune competence, since injection of live bacteria provides an integrative view of the immune system activation of the individual (Charles & Killian, 2015), and different parameters linked to antimicrobial immune response in insects are positively correlated (Gillespie et al. 1997; Schmid-Hempel 2005). Furthermore, *E. coli* is not naturally found in *A. mellifera* (Cini et al. 2020), so we could therefore exclude its presence in our workers prior to artificial infection. Bacterial cultures of *E. coli* tetracycline-resistant strain XL1-Blue were grown in Luria-Bertani (LB) complex medium added with 10 μg/mL tetracycline overnight at 37 °C in a shaking incubator. After centrifugation, bacteria were washed twice and then resuspended in phosphate buffered saline (PBS). The approximate number of bacterial cells in the solution was measured as optical density with an Eppendorf BioPhotometer® 6131, then cells were diluted to ∼ 1.5 × 10^8^ cells/mL in PBS. Bees were injected with 1 μL of inoculum, containing ∼ 1.5 × 10^5^ cells, between the 2^nd^ and 3^rd^ tergite with a Hamilton^TM^ micro-syringe (Cappa et al. 2020). After injection, bees were separated according to category into 120 ml Plexiglass cages provided with ad libitum 50% sugar solution as food and maintained under controlled conditions (∼ 30 °C; 55% RH). Twenty-four hours later workers were dissected in a plate on ice to remove the sting apparatus in order to avoid a reduction in bacteria viability due to antimicrobial activity of venom compounds (Baracchi et al. 2012). Each worker was then inserted into a 1.5 mL Eppendorf containing 1 mL of sterile PBS and the bee body was manually homogenized for 1 minute using a sterile metal pestle. Afterwards, 0.1 mL of undiluted and serially diluted PBS suspensions (dilutions 10^−1^-10^−4^) of each sample were plated on LB solid medium added with tetracycline (10 μg/mL) and incubated overnight at 37 °C. The following day, the colonies grown on the plate surface were counted and the viable bacterial count was expressed as Colony Forming Units (CFUs) per mL. At least three bees per colony were injected with 1 μL of PBS, homogenized and plated following the same procedure of *E. coli-* infected workers, to ensure absence of other bacterial strains capable of growing on our LB agar plates added with tetracycline. A total of 258 bee were infected with *E. coli* and plated (C-PS = 23, C-PMMA = 23, C-MIX = 15, PS-L = 25, PS-M = 24, PS-H = 23, PMMAL = 23, PMMAM = 23, PMMAH = 23, MIXL = 20, MIXM = 20, MIXH = 16). Bacterial challenge data as raw number of CFU/mL were Log-transformed for the statistical analyses.

### Cuticular Analysis

To analyse the profiles of cuticular hydrocarbons (CHCs), we collected a total of 240 bees from two colonies, with 120 bees from each colony used in the behavioural test described below. These bees were housed in cages with 30 individuals per cage. Each cage was randomly assigned to either the control or experimental treatment group (as described above). After a 3-day exposure period, the bees were killed by freezing at −20°C. Epicuticular compounds were extracted from each bee by washing for 10 min in 400 μL pentane in 2 mL vials. The samples were then allowed to evaporate completely and resuspended in 100 μL pentane. The samples were then vortexed for 10 sec. The resuspended extract was transferred to a 250 μL glass conical insert, dried under a nitrogen stream and suspended in 80 μL heptane containing 70 ng/μL hexadecan-1-ol (C_16_OH as an internal standard. We injected 1 μL of solution into Hewlett Packard (Palo Alto, CA, USA) 5890A gas chromatograph (GC) coupled to an HP 5970 mass selective detector (using a 70-eV electronic ionization source). A fused ZB-WAX-PLUS (Zebron) silica capillary column (60 m × 0.25 mm × 0.25 mm) was installed in the GC. The injector port and transfer line temperatures were set at 200 °C and the carrier gas was helium (at 20 PSI head pressure). The temperature protocol was from 50 °C to 320 °C at a rate of 10 °C/min, and the final temperature was held for 5 min. Injections were performed in splitless mode (1 min purge valve off). Data acquisition and analysis were performed using the Agilent Masshunter Workstation software (version B.07.00). Compounds were identified on the basis of their retention time, mass spectra and equivalent chain length. Mass spectra, with their corresponding molecular weights, were compared with electronic mass spectra libraries (NIST MS Search v.2.0). A total of 45 peaks corresponding to different compounds were included in the data set. The peak areas of the chromatogram for each bee were normalised to the area of the internal standard (C_16_OH). For the identification and analysis of CHCs, peaks quantified in less than 10% of the samples were excluded, except for those quantified in only one of the treatment groups. A total of 38 compounds were included in the statistical analysis. Due to lack of data or excessive background noise in the chromatograms, four bees from the control group and five bees exposed to the low concentration of PS (PSL) were not included in the analysis. A total of 231 individuals (C = 56, PSL = 15, PSM = 20, PSH = 20, PMMAL = 20, PMMAM = 20, PMMAH = 20, MIXL = 20, MIXM = 20, MIXH = 20) were included in the statistical analysis.

### Behavioural Assay

To investigate the effect of MPs on recognition mechanisms among nestmates, and in particular the ability of guard bees at the hive entrance to detect exposed bees, we collected 360 bees from three colonies, with 120 bees from each colony. These bees were then caged in groups of 20 and randomly assigned to either control or experimental groups. Bees in the control group were fed a 50% (w/w) sucrose solution, while those in the experimental groups were fed PMMA, PS or MIX, each at different concentrations (see above). After 3 days of exposure, the bees were returned to the hive entrance of their respective colonies. Each experimental bee was briefly placed in an ice-cooled box for one minute prior to release to calm the bee and prevent immediate flight upon release. At the start of the test, each focal bee was gently released onto the corner of the hive’s landing board, allowing free movement to interact with other bees, enter the hive or take flight. The focal bee was recorded for 3 min after the first interaction with any bee on the landing board or until it entered the hive. If the focal bee flew away before any interaction occurred, that assay was disregarded. Videos were blind watched at half speed to record the number and the duration of each interaction between focal bees and bees on the landing board. The average number of bees at the entrance was calculated by counting the number of bees on the flight board at the beginning and at the end of each presentation (Cappa et al. 2014). Each interaction was categorised based on a predefined list of behaviours, which were grouped into two main categories: affiliative or aggressive acts. Affiliative behaviours included interactions such as antennal inspection, allo-grooming and trophallaxis. Aggressive behaviours, on the other hand, included biting, darting, pulling, and removing of the focal bee from the hive entrance.

### Statistical Analysis

Linear Mixed Models (LMM) were used to examine the dependence of dietary intake on groups (Raudenbush and Bryk, 2002). Mixed-effects Cox regression (Fox and Weisberg, 2002; Lin and Zelterman, 2002) was used to estimate the effect of groups on survival rates. Non-parametric models for repeated-measures ANOVA, Kruskal-Wallis’s test (Kruskal & Wallis, 1952), were used to analyse immune competence data and individual behaviours derived from the behavioural assay. Behavioural data were also explored using Principal Component Analysis (PCA) to identify potential associations between all measured variables. All behaviours were normalised according to the number of bees at the hive entrance at the beginning and end of each test. Partial Least Squares Discriminant Analysis (PLS-DA), implemented in open-source software MetaboAnalyst, was used to assess sample separation, and determine the specific compounds contributing to class separation in the cuticular hydrocarbons (CHCs) of bees. Prior to analysis, data were log10 transformed to better approximate normality and homogeneity of variances. Mixed-effects models for repeated measures ANOVA were used to assess differences between groups on a peak-by-peak basis. All statistical analyses were performed in R 4.3.1 (R Core Team, 2021).

## Results

### Survival Assay

Across all groups of confined bees, food consumption consistently decreased during the test period (LMM, *day*, PMMA: *p* < 0.0001; PS: *p* < 0.0001; MIX: *p* < 0.0001, Suppl. Figure SM1). Chronic oral treatment with PS, PMMA, or the MIX consistently showed no discernible effect on the per capita daily food consumption, irrespective of the concentration applied (LMM, *treatment*, PMMA: *p* = 0.36; PS: *p* = 0.9; MIX: *p* = 0.72, Suppl. Figure SM1). Survival curves are shown in Figure 1. The estimated mixed effects Cox regression indicated a difference in survival rates between the MIX group and its control group (Log-Rank test: χ^2^ = 10.43, df = 3, *p* = 0.015), whereas no such difference was observed between the PMMA and PS groups and their respective controls (Log-Rank test: PMMA: χ^2^ = 0.72, df = 3, *p* = 0.87; PS: χ^2^ = 2.72, df = 3, *p* = 0.44). Accordingly, PMMA and PS treatments show no negative correlation with survival time (PMMA: H: *p* = 0.40; M: *p* = 0.54; L: *p* = 0.58; PS: H: *p* = 0.26; M: *p* = 0.83; L: *p* = 0.45). Bees exposed to a combination of the two MPs at high and medium concentrations experienced faster mortality compared to the control bees (MIX: H: *p* = 0.015; M: *p* = 0.005). However, this pattern was not observed at lower concentrations (*p* = 0.32).

**Figure 1:**
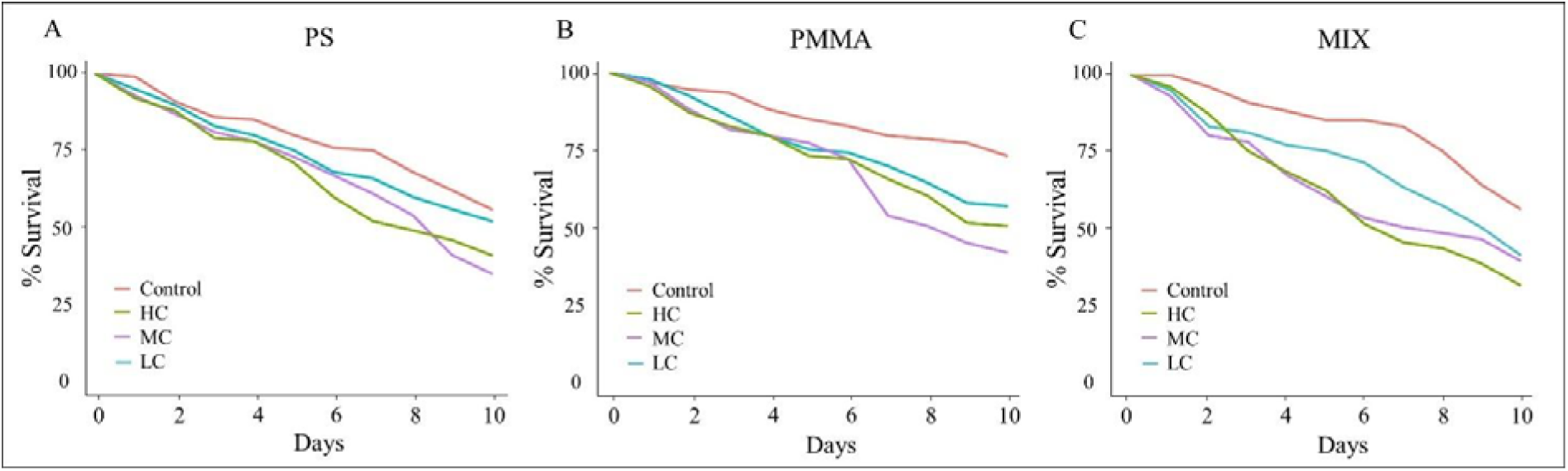
Cumulative survival of MPs exposed to either PS (A), PMMA (B) or MIX (C). Survival rate (y-axis) of bees exposed to MPs at different concentrations (High (H) = green, Low (L) = blue, Medium (M) = purple) and control bees (C = red) kept in cages over 10 days (x-axis). Analysis showed a difference in survival rates between the MIX group and its control group (*p* = 0.015), whereas no such difference was observed between the PS and PMMA groups and their respective controls (*p* = 0.87 and *p* = 0.44). Bees exposed to MIX at high and medium concentrations experienced faster mortality compared to the control bees (H: *p* = 0.015; M: *p* = 0.005), whereas no differences were observed at low concentration (*p* = 0.32).

### Immunocompetence Assay

As regards exposure to PS, the analyses showed a significant difference among the groups exposed to different concentrations (χ^2^ = 38.295, df = 3, *p* < 0.001) (Figure 2A). In particular, a higher CFUs value, corresponding to a reduced bacterial clearance, was found in the bees exposed to the highest concentration (H) of PS in comparison with the control group (F = 5.36, *p* < 0.001). No significant difference emerged between the control bees and those exposed to the other concentrations (C - PS- L: F = 0.95, *p* = 0.34; C - PS-M: F = 0.246, *p* = 0.81).

**Figure 2:**
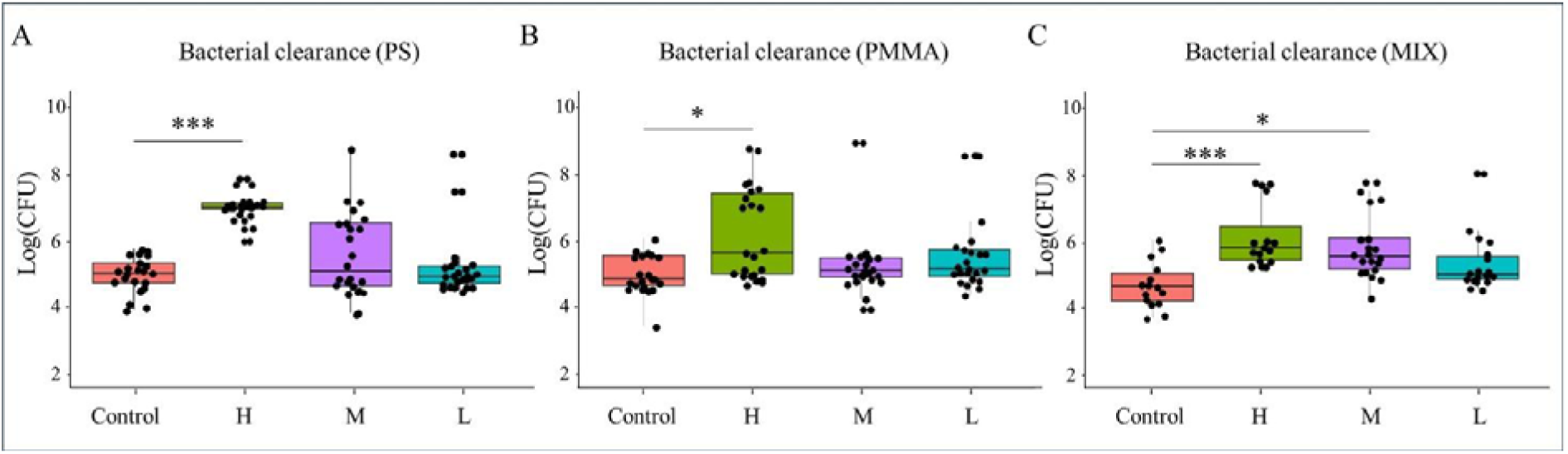
Bacterial count (expressed as CFUs) in bees exposed to either PS (A), PMMA (B) or MIX (C). Box plots show CFUs (y-axis) in bees exposed to different concentrations of MPs (High (H) = green, Low (L) = blue, Medium (M) = purple) and control bees (C = red). Significantly higher bacterial counts were observed in bees exposed to H-PS and H-PMMA compared to the corresponding control group. Bees exposed to H-MIX and M-MIX had higher bacterial counts than controls. Notably, no significant differences were found between control bees and those exposed to lower concentrations in any of the groups. The dots represent single bees. (*) = p < 0.05; (**) = p < 0.01; (***) = p < 0.001.

Even for bees exposed to different concentrations of PMMA, the analysis showed a significant difference among the groups (χ^2^ = 11.187, df = 3, *p* = 0.01, Figure 2B). Specifically, a higher bacterial count was once more observed in the bees exposed to the highest concentration (H) of PMMA compared to the control group (F = 3.21, *p* = 0.008). No significant differences were observed among the control bees and those exposed to the other concentrations (C – PMMA-L: F = 1.08, *p* = 0.28; C – PMMA-M: F = 1.98, *p* = 1). Finally, also for bees exposed to the MIX at different concentrations, the analyses showed a significant difference among groups (χ^2^ = 23.281, df = 3, *p* < 0.001, Figure 2C). Interestingly, higher CFUs values were found not only in bees exposed to the highest concentration (H) of MIX compared to the control group (F = 4.42, *p* < 0.001), but also in the group of bees exposed to the field-realistic medium concentration (M) of MIX compared to controls (F = 3.57, *p* = 0.002). No significant difference emerged among control bees and those exposed to the lowest concentration.

### Chemical analysis

The 38 identified compounds included in the analyses belonged to four different classes: alkanes (N = 13), alkenes (N = 12), methyl-branched hydrocarbons (N = 2), esters between fatty acids and fatty alcohol (N = 10), plus one unidentified compound. The compounds identified in the cuticular profiles were very similar to those found in bees from the same area in a previous study (see Cappa et al., 2019). To evaluate the separation between samples and predefined groups, we performed a PLS-DA which allowed us to visualize the general pattern of similarity between individuals exposed to PS (Figure 3A), PMMA (Figure 3B), MIX (Figure 3C) and control bees. For all MPs treatments, exposed bees showed good separation from controls.

**Figure 3:**
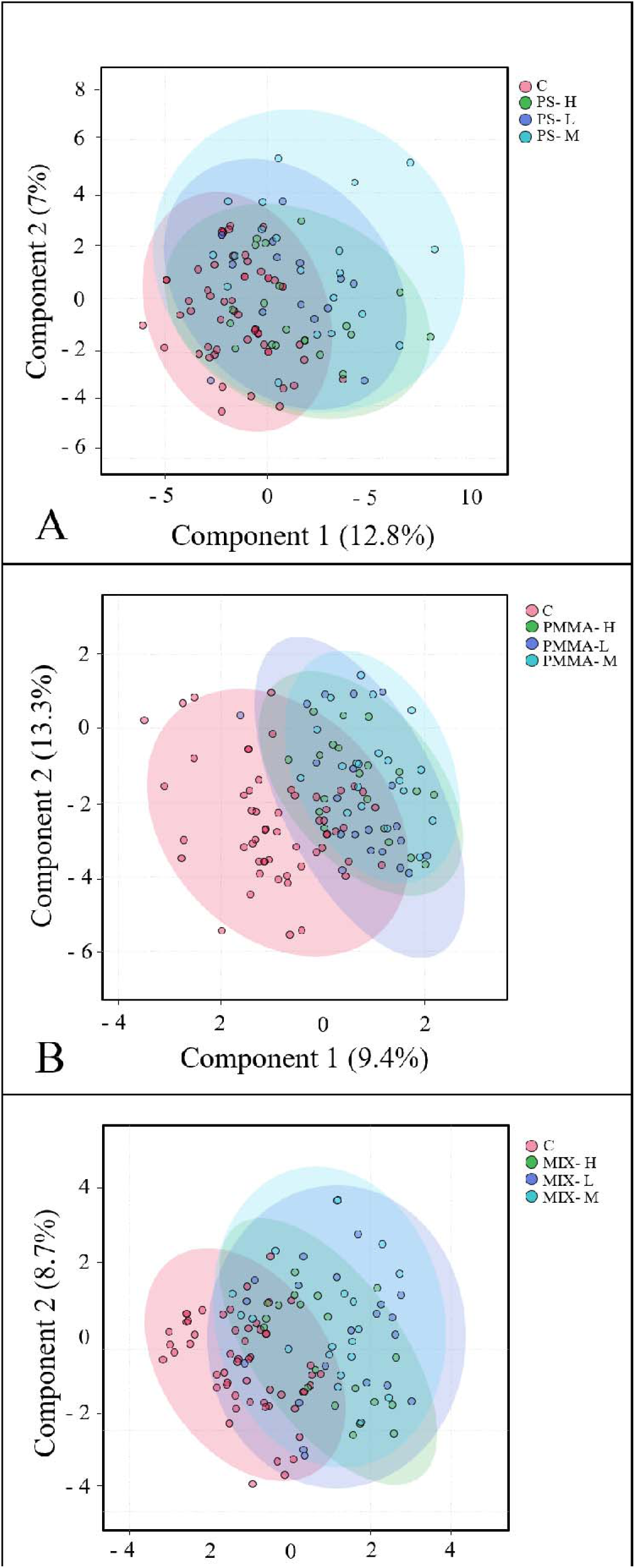
Partial Least Squares-Discriminant Analysis of CHCs profile in honey bees exposed to PS (A), PMMA (B) or MIX (C). Scatterplot for the first component (x-axis) and second component (y-axis) of a Partial Least Squares Discriminant Analysis on bees exposed to different MPs concentrations (High (H) = green, Low (L) = blue, Medium (M) = purple) and control bees (C = red). Each dot represents a single bee. For all MPs treatments, exposed bees showed good separation from controls.

To better understand which compounds allowed class separation, individual hydrocarbons were subjected to ANOVA across all groups. Table 1 shows the amount of cuticular hydrocarbons in bees treated with MPs at different concentrations (L, M, H) compared to control bees. Comparing Ps-treated bees with control bees, we found significant differences in 8 compounds: three alkanes (C29, *p* < 0.001; C30, *p* = 0.015; C31, *p* = 0.006), three alkenes (C29:1, *p* = 0.26; C31:2, *p* = 0.005; C31:1b, *p* = 0.001) and two fatty acid esters (Oleic acid ester 3, p = 0.025; Palmitic acid ester 1, *p* = 0.03). Post hoc analyses showed no differences between bees treated with the lowest concentration of PS (PS-L) and control bees. Bees exposed to the medium concentration (PS-M) showed lower amounts than controls of C29:1 (*p* = 0.04), C30 (*p* = 0.019) and higher amounts of C31:2 (*p* = 0.044). Bees exposed to the highest concentration (PS-H) showed lower amounts of C29 (*p* < 0.001), C31:1b (*p* < 0.001), C31 (*p* = 0.004), an Oleic acid ester 3 (*p* = 0.017) and higher amounts of a Palmitic acid ester 1 (*p* = 0.04). For detailed results see Suppl. Table SM1. Comparing bees treated with PMMA to controls, we found significant differences in 18 compounds: four alkanes (C21, *p* < 0.001; C22, *p* = 0.01; C26, *p* = 0.02; C27, *p* < 0.001), nine alkenes (C23:1a, *p* < 0.001; C23:1b, *p* = 0.01; C24:1, *p* < 0.001 ; C25:1a, *p* = 0.01; C25:1b, *p* = 0.007; C27:1a, *p* < 0.001 ; C27:1b, *p* < 0.001; C29:1, *p* < 0.001 ; C31:2, *p* = 0.007), one methyl of C27 (n-methylC27, *p* = 0.01) three fatty acid esters (Oleic acid ester 1, *p* = 0.02; Oleic acid ester 5, *p* < 0.001, Palmitic acid ester 2, *p* < 0.001) and an undefined compound (*p* < 0.001). Post hoc analyses showed that bees exposed to PMMA at all three concentrations had higher amounts of C24:1 (PMMA-L - C, *p* = 0.03; PMMA-M - C, *p* = 0.001; PMMA-H - C *p* = 0.002) and lower amounts of an oleic acid ester (Oleic acid ester 5, PMMA-L - C, *p* = 0.001; PMMA-M - C, *p* = 0.001; PMMA-H - C *p* = 0.001).

**Table 1:** Amount of cuticular hydrocarbons in bees treated with PS, PMMA and MIX at different concentrations (L, M and H) compared to control bees. The table gives *p-values* from the ANOVA performed on each chemical compound and indicates the concentrations found to be different from controls in post hoc analysis. When PS-treated bees were compared with control bees, significant differences were observed for 8 compounds, including three alkanes, three alkenes and two fatty acid esters. Bees exposed to PMMA showed distinct concentrations of 18 compounds compared to controls, including four alkanes, nine alkenes, one methyl of C27, three fatty acid esters and one undefined compound. Analysis of bees exposed to MIX showed quantitative differences from controls in 17 compounds, including seven alkanes, six alkenes and three fatty acid esters. Further detailed results are presented in Suppl. Tables SM 1, 2 and 3.

| Compound | PS |  | PMMA |  | MIX |  |
| --- | --- | --- | --- | --- | --- | --- |
|  | *p* | Post-hoc | *p* | Post-hoc | *p* | Post-hoc |
| C<sub>19</sub> | 0.853 |  | 0.232 |  | 0.1 |  |
| Octadecenoic Acid | 0.307 |  | 0.1 |  | 0.4 |  |
| C<sub>21</sub> | 0.741 |  | < 0.001 *** | L | 0.035 * |  |
| C<sub>22</sub> | 0.465 |  | 0.01 * | M | 0.01 * | H |
| 11-Eicosen-1-ol | 0.851 |  | 0.02 |  | 0.002 ** | H |
| C<sub>23:1a</sub> | 0.195 |  | 0.01 * | M | 0.4 |  |
| C<sub>23:1b</sub> | 0.628 |  | 0.02 * | M | 0.03 * | M |
| C<sub>23</sub> | 0.286 |  | 0.119 |  | 0.9 |  |
| C<sub>24:1</sub> | 0.07 |  | < 0.001 *** | L, M, H | 0.5 |  |
| C<sub>24</sub> | 0.215 |  | 0.066 |  | 0.35 |  |
| C<sub>25:1a</sub> | 0.393 |  | 0.01 ** | M | 0.88 |  |
| C<sub>25:1b</sub> | 0.396 |  | 0.007 ** | M | 0.8 |  |
| C<sub>25</sub> | 0.256 |  | 0.06 |  | 0.04 * |  |
| C<sub>26</sub> | 0.408 |  | 0.02 * | H | < 0.001 *** | H |
| C<sub>27:1a</sub> | 0.087 |  | < 0.001 *** | L, H | 0.002 ** | L |
| C<sub>27:1b</sub> | 0.8 |  | < 0.001 *** | M | < 0.001 *** | L, H |
| C<sub>27</sub> | 0.057 |  | < 0.001 *** | L, H | < 0.001 *** | L, H |
| *n*-meC<sub>27</sub> | 0.052 |  | 0.01 ** |  | 0.421 |  |
| C<sub>28</sub> | 0.21 |  | 0.136 |  | 0.8 |  |
| C<sub>29:1</sub> | 0.026 * | M | < 0.001 *** | H | 0.007 ** |  |
| C<sub>29</sub> | < 0.001 *** | H | 0.067 |  | < 0.001 *** | L, H |
| *n*-meC<sub>29</sub> | 0.478 |  | 0.28 |  | 0.25 |  |
| C<sub>30</sub> | 0.015 * | M | 0.4 |  | 0.08 |  |
| C<sub>31:2</sub> | 0.005 * | M | 0.007 ** | H | 0.07 |  |
| C<sub>31:1a</sub> | 0.118 |  | 0.7 |  | 0.1 |  |
| C<sub>31:1b</sub> | < 0.001 *** | H | 0.68 |  | 0.015 * | L |
| C<sub>31</sub> | 0.006 ** | H | 0.13 |  | < 0.001 *** | L |
| Oleic Acid ester 1 | 0.498 |  | 0.02 * | H | 0.06 |  |
| Oleic Acid ester 2 | 0.427 |  | 0.1 |  | 0.027 * | L |
| Oleic Acid ester 3 | 0.025 * | H | 0.35 |  | 0.1 |  |
| C<sub>33:1</sub> | 0.07 |  | 0.08 |  | 0.002 ** | L |
| C<sub>33</sub> | 0.066 |  | 0.1 |  | 0.7 |  |
| Oleic Acid ester 4 | 0.536 |  | 0.06 |  | 0.45 |  |
| Undefined | 0.949 |  | < 0.001 *** | M, H | 0.4 |  |
| Oleic Acid ester 5 | 0.379 |  | < 0.001 *** | L, M, H | 0.01 ** | H |
| Palmitic Acid ester 1 | 0.03 * | H | 0.16 |  | < 0.001 *** | L, H |
| Palmitic Acid ester 2 | 0.984 |  | < 0.001 *** | M, H | 0.07 |  |
| Palmitic Acid ester 3 | 0.103 |  | 0.4 |  | 0.5 |  |

Bees exposed to the M and H concentrations also showed higher amounts of a palmitic acid ester 2 (PMMA-M - C, *p* < 0.001; PMMA-H – C, *p* = 0.001). Overall, PMMA-L showed quantitative differences with control in 5 compounds, PMMA-M in 9, while PMMA-H in 8 compounds. For detailed results see Suppl. Table SM2. Regarding profiles of bees exposed to the MIX at different concentrations (L, M, H), analyses revealed quantitative differences with controls in 17 compounds: seven alkanes (C21, F *p* = 0.035; C22, *p* = 0.01; C25, *p* = 0.04; C26, *p* < 0.001; C27, *p* < 0.001; C29, *p* < 0.001; C31, *p* < 0.001), six alkenes (C23:1b, *p* = 0.03; C27:1a, *p* = 0.002; C27:1b, *p* < 0.001; C29:1, *p* = 0.007; C31:1b, *p* = 0.015; C33:1, F *p* = 0.002) and three fatty acid esters (Oleic acid ester 2, *p* = 0.27; Oleic acid ester 5, *p* = 0.01; Palmitic acid ester 1, *p* < 0.001; 11-eicosen-1-ol, *p* = 0.002). Post hoc analyses showed no differences between bees treated with the medium concentration (MIX-M) and control bees. Bees exposed to the lowest concentration showed quantitative differences with controls in 10 compounds, while the highest one in 7. For results in detail see Suppl. Table SM3.

### Behavioural Assay

To evaluate possible associations between all measured variables, we performed a PCA on control bees and bees exposed to PS, PMMA, and the MIX. The PCAs extracted 10 PCs explaining 100% of the variance. Figure 4A shows the first two components from the PCA performed on bees exposed to PS at different concentrations (L, M, H) and control bees. PC1 explained the 31.4% of the variance, while PC2 explained 21.3%. The most important variables in PC1 were affiliative acts (18%), and the number and the duration of antennations (17.5% and 16% respectively), while the most important ones in PC2 were aggressive acts (20.5%), bites events (18.6%), and dragging events (12.8%) (details in Suppl. Table SM4). Figure 4B shows PC1 (34.1%) and PC2 (16.8%) from PCA performed on PMMA exposed individuals and control bees. The most important variables in PC1 were affiliative acts (15%), and the number and the duration of antennations (14.6% and 15.2%, respectively), while the most important ones in PC2 were the number and the duration of grooming (40.5% and 41.4 % respectively), and the number and the duration of trophallaxis (5.8% and 5.9%, respectively) (details in Suppl. Table SM4). Figure 4C shows PC1 (31%) and PC2 (16.6%) from PCA performed on MIX exposed bees. Affiliative acts (17.3%), and the number and duration of antennations (15.3% and 17.5%, respectively) were the most important variables in PC1, while the number and duration of trophallaxis (33.4% and 32.3%, respectively) and those of grooming (9.2% and 9.1%, respectively) were the most important in PC2 (details in Suppl. Table SM4). Overall, these results suggest that the experimental and control groups appeared to largely overlap.

**Figure 4:**
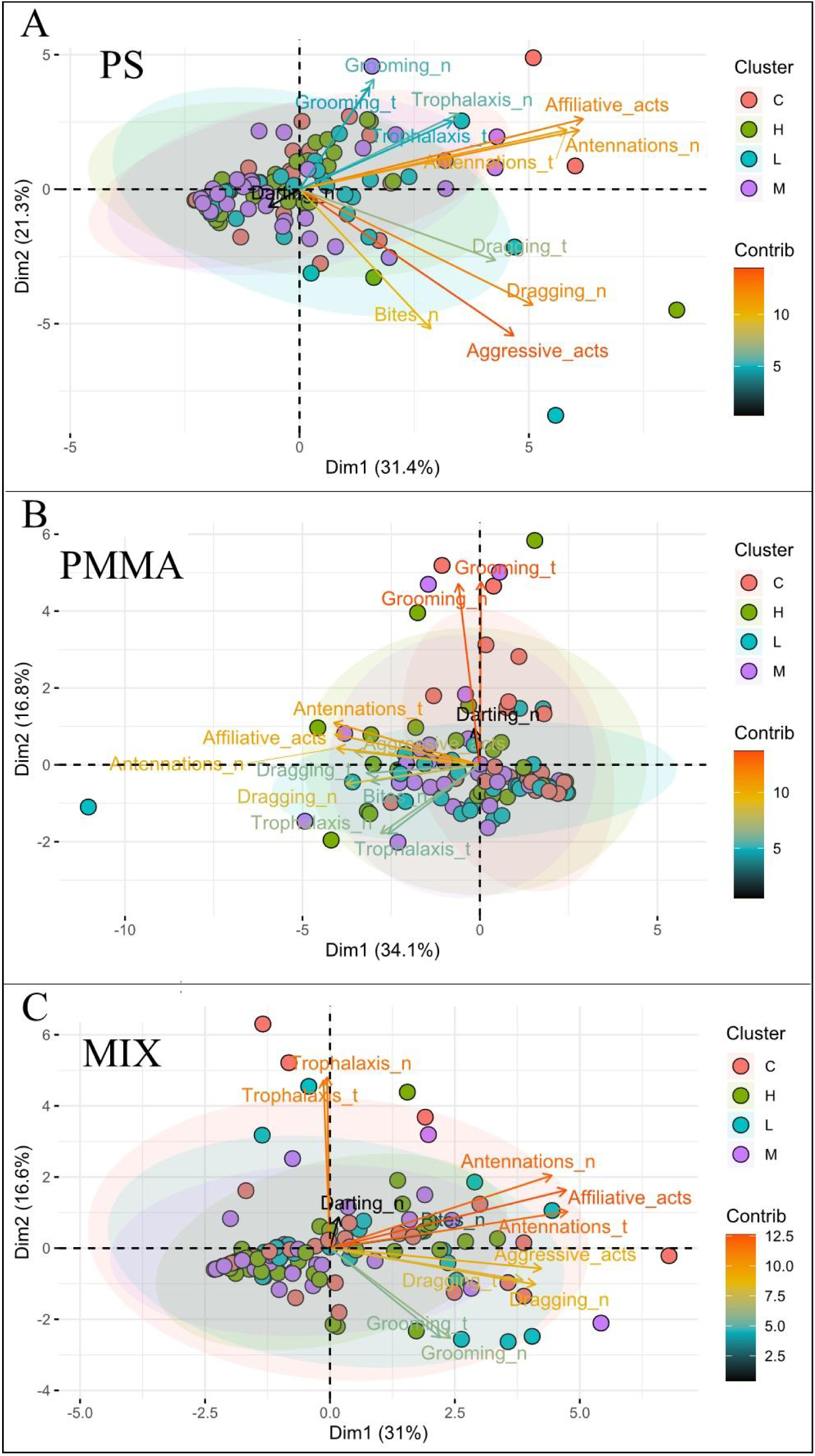
Principal Component Analysis on Control bees and bees exposed to MPs. Scatterplot for the first two components of a PCA on bees exposed to different concentrations of MPs (High (H) = green, Low (L) = blue, Medium (M) = purple) and control bees (C = red). **A**: PCA output from bees exposed to PS and control bees. **B**: PCA output from bees exposed to PMMA and control bees. **C**: PCA output from bees exposed to MIX and control bees.

Kruskal-Wallis tests were performed on each behaviour to properly assess differences between groups. Figure 5 shows the average of affiliative and aggressive acts towards control bees and those exposed to MPs. Comparison between control bees and those exposed to PS or PMMA showed no differences in either affiliative (PS: χ^2^ = 1.51, df = 3, *p* = 0.68; PMMA: χ^2^= 5.61, df = 3, *p* = 0.13) or aggressive acts (PS: χ^2^ = 5.82, df = 3, *p* = 0.12; PMMA: χ^2^ = 2.14, df = 3, *p* = 0.54). Detailed analysis of individual behaviours also revealed no significant differences (see Suppl. Table SM5 and 6). Similarly, the analysis of bees exposed to the MIX showed no significant differences compared to control bees, neither for affiliative acts (χ^2^ = 1.05, df = 3, *p* = 0.79) and aggressive acts (χ^2^ = 1.88, df = 3, *p* = 0.60) nor for individual behaviours (see Suppl. Table SM7).

**Figure 5:**
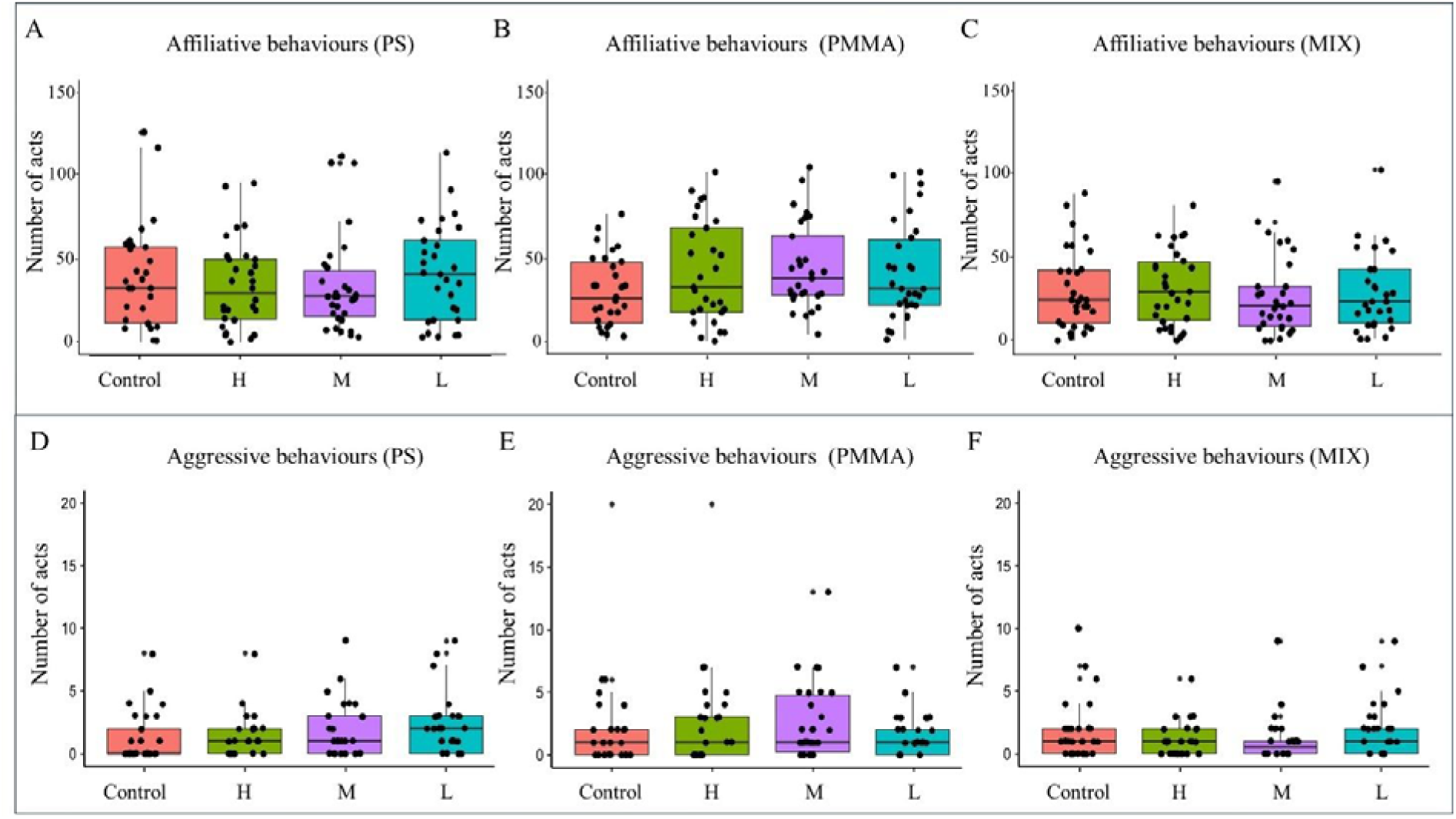
Affiliative and aggressive acts toward control bees and bees exposed to MPs. Box plots showing average affiliative and aggressive acts (Y-axis) towards control bees (C = red) and those exposed to different concentrations of MPs -High (H) = green, Low (L) = blue, Medium (M) = purple (X-axis). Panels A, B, and C display combined data for PS, PMMA, and MIX exposure, respectively. No significant differences were observed in either affiliative or aggressive acts between control bees and those exposed to the respective MPs or their combinations.

## Discussion

Our study pioneers the investigation of the effects of PS, PMMA, and their combination on the survival, immune system, and colony integrity of a social insect, and is the first investigation of its kind in any organism. We found a synergistic effect of the two MPs on survival and the immune system. Furthermore, both MPs, whether administered singly or in combination, altered the cuticular profile of exposed bees. However, this alteration did not result in increased aggressive or inspection behaviours toward exposed bees at the hive entrance, which were able to enter the colony as unexposed individuals.

Specifically, survival tests showed a negative effect on the survival rate of bees exposed to MPs administered in combination, whereas no such difference was observed between the PMMA and PS groups and their respective controls. Interestingly, the reduction in survival rates occurred already at the medium concentration of the mixture (5 mg/L). This concentration falls in the field-realistic range (Chae et al., 2018; Eltemsah and Bohn, 2019; Wang et al., 2021, making these results even more alarming. In fact, simultaneous exposure to field concentrations of these two MPs during foraging could jeopardize the survival of honey bees. This concern is further magnified by the knowledge that free-living organisms are typically exposed to a combination of contaminants, as highlighted by previous research (Siviter et al., 2021; Al Naggar et al., 2022). Our study, which is the first to demonstrate a synergistic effect on honey bee survival due to combined exposure to PS and PMMA, highlights the tangible threat these MPs pose to bees, and, most importantly, the risk extends to concentrations potentially encountered by foragers in the field.

These findings are supported by the results of our immunocompetence assay. Even when administered individually, both PS and PMMA resulted in a weakened immune response in exposed workers. Notably, this impairment occurred only at the highest concentration of MPs (50 mg/L), a level well above the field-realistic range of variability found in natural environment. In contrast, combined exposure to PS and PMMA impaired the immune competence of bees even at the field-realistic concentration of 5 mg/L, suggesting potential synergistic interactions between the two MPs that could plausibly occur in an ecological context. Our immunocompetence assay results align with recent studies showing a correlation between MP intake and activation of genes associated with immune response and detoxification (Deng et al., 2021; K. Wang et al., 2022). An impaired immune system is bound to have a significant impact on various aspects of bee life. For example, non-pathogenic activation of the immune system has been shown to alter learning and memory in honey bees and bumble bees (Mallon et al., 2003; Alghamdi et al., 2008). Furthermore, exposure to MPs may exacerbate susceptibility to additional stressors, including microbial infections, parasites, and various classes of pollutants, by compromising the bees’ immune defences. This complex interplay, coupled with the observed adverse effects on survival, broadens the risks associated with MPs and potentially amplifies the overall vulnerability of honey bee populations to these and other environmental stressors.

Activation of the immune system has been shown to induce changes in the chemical profile (CHCs) of honey bees (Baracchi et al., 2012; Richard et al., 2008, 2012; Cappa et al. 2016). Our chemical analyses showed that both MPs induced a change in the cuticular profile of bees. In general, exposure to MPs resulted in a reduction in the abundance of several cuticular compounds, although we did not observe a specific pattern of alteration with respect to the different MPs or their increasing concentrations. The reduction in CHCs in exposed foragers may explain the apparently unexpected results of our behavioural assays. Hive entry guards did not show increased inspection or aggressive behaviour towards exposed foragers compared to unexposed foragers. Previous studies in social insects have found a correlation between aggressive behaviour and higher concentrations of CHCs (Cini et al., 2009; Ichinose & Lenoir, 2010; Cappa et al. 2016). In honey bees, Cappa et al. (2014) demonstrated that exploratory and aggressive responses towards non-nestmates increased with increasing cue concentration, following a threshold mechanism. Moreover, Cappa et al. (2019) observed that exposure to the entomopathogenic fungus *Beauveria baussiana* caused a reduction in specific CHCs in honey bee foragers, leading to higher hive acceptance of non-nestmates compared to those with an unaltered profile. Consistent with these findings, our experiment showed that exposure to MPs tended to reduce levels of several CHCs, potentially mitigating an aggressive response in guard bees. The failure to recognise bees exposed to MPs could potentially facilitate the penetration and spread of these contaminants within the hive, a hypothesis consistent with previous studies detecting MPs in the hive and in various bee products such as honey and wax (Alma et al., 2023; Diaz-Basantes et al., 2020). In addition, the observed immunosuppressive effects of MPs exposure could increase the entry and spread of pathogens and parasites carried by immunodeficient bees within the colony. Finally, a general reduction in the amount of CHCs in foragers exposed to MPs could increase the likelihood of acceptance of exposed non-nestmates into the colony (Cini et al., 2009; Ichinose & Lenoir, 2010; Cappa et al., 2014, 2019). This possibility must be urgently investigated in the future, as it is well known that the entry of foreign individuals, who may steal resources or introduce new diseases, poses a serious threat to the overall integrity and survival of the colony.

In conclusion, our study demonstrates how exposure to PS and PMMA significantly affects honey bee health at different levels. For the first time, we found that these two MPs appear to interfere synergistically with bee survival and immunity when exposed simultaneously. The occurrence of this synergistic interaction is of particular concern as it is manifested even at field-realistic concentrations. We also observed an alteration in the cuticular profile of bees exposed to MPs. Despite these chemical changes, nest guards appear unable to detect exposed foragers, allowing them to enter the colony undisturbed. This lack of an effective detection system could potentially facilitate the spread of MPs and other contaminants or pathogens within the hive, posing serious risks to the entire colony. The recognition of these implications underlines the urgency of stepping up research efforts to understand the threat posed to pollinators by plastic pollution, a major challenge caused by human activity.

## Author Contributions

D.B. planned and designed the study, F.F., E.P., L.B. performed behavioural experiments. E.P. and F.C. performed immunological experiments. F.F., E.P., L.B. and F.C. performed chemical analysis. D.B. and F.F. performed statistical analysis. D.B. provided funding for the research. F.F. and D.B. wrote the first draft of the manuscript. All authors contributed to writing the final version of the manuscript and gave final approval for publication.

## Acknowledgements

We acknowledge the assistance of Giuseppe Pieraccini and Giulia Ricciardi for chemical analysis at the Mass Spectrometry Center (CISM) and of Maria Theocharidou for behavioural experiments.

