## Supplementary Materials for "Unravelling the microplastic menace: different polymers work in synergy to increase bee vulnerability"

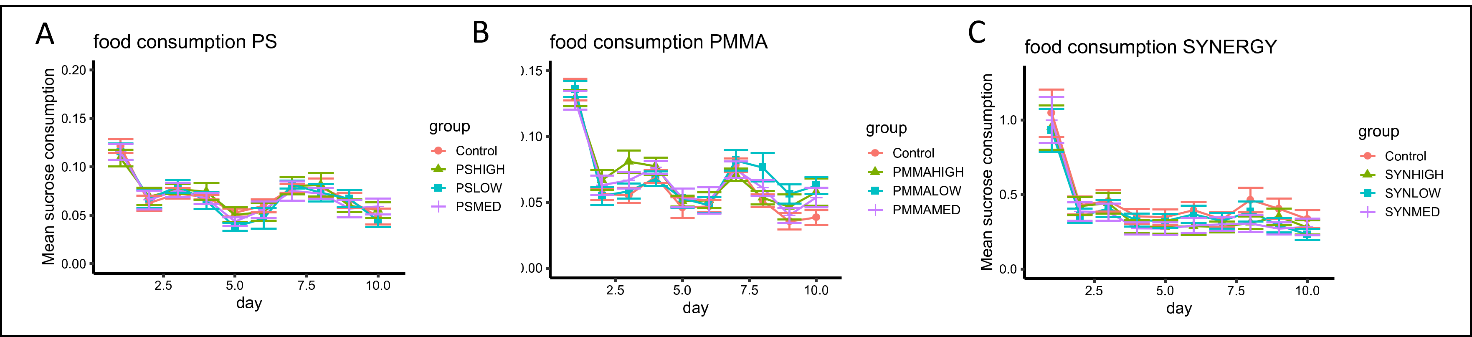

**Figure SM1.** **Food consumption in bees exposed to either PS (A), PMMA (B) or MIX (C).** Across all groups, food consumption consistently decreased during the test period (PS: *p* < 0.0001; PMMA: *p* < 0.0001; MIX: *p* < 0.0001). No discernible effect on the per capita daily food consumption was observed regardless of treatment (PS: *p* = 0.9; PMMA: *p* = 0.36; PS: *p* = 0.9; MIX: *p* = 0.72).

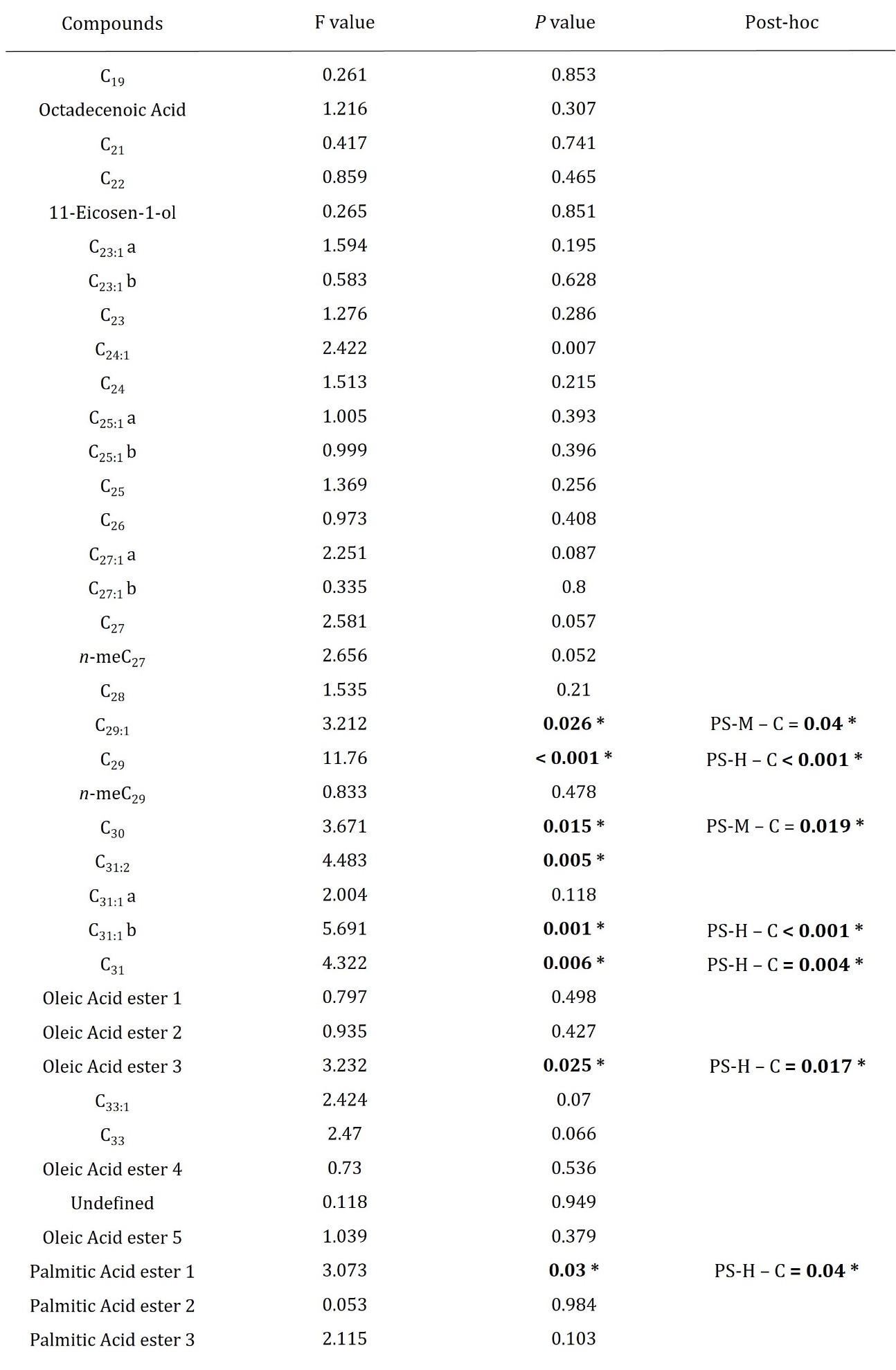

**Table SM1: Amount of cuticular hydrocarbons in bees treated with PS (L, M and H) compared to control bees.** Table gives F values and *p* values from ANOVA performed on each compound and *p* value from post-hoc analysis across all groups.

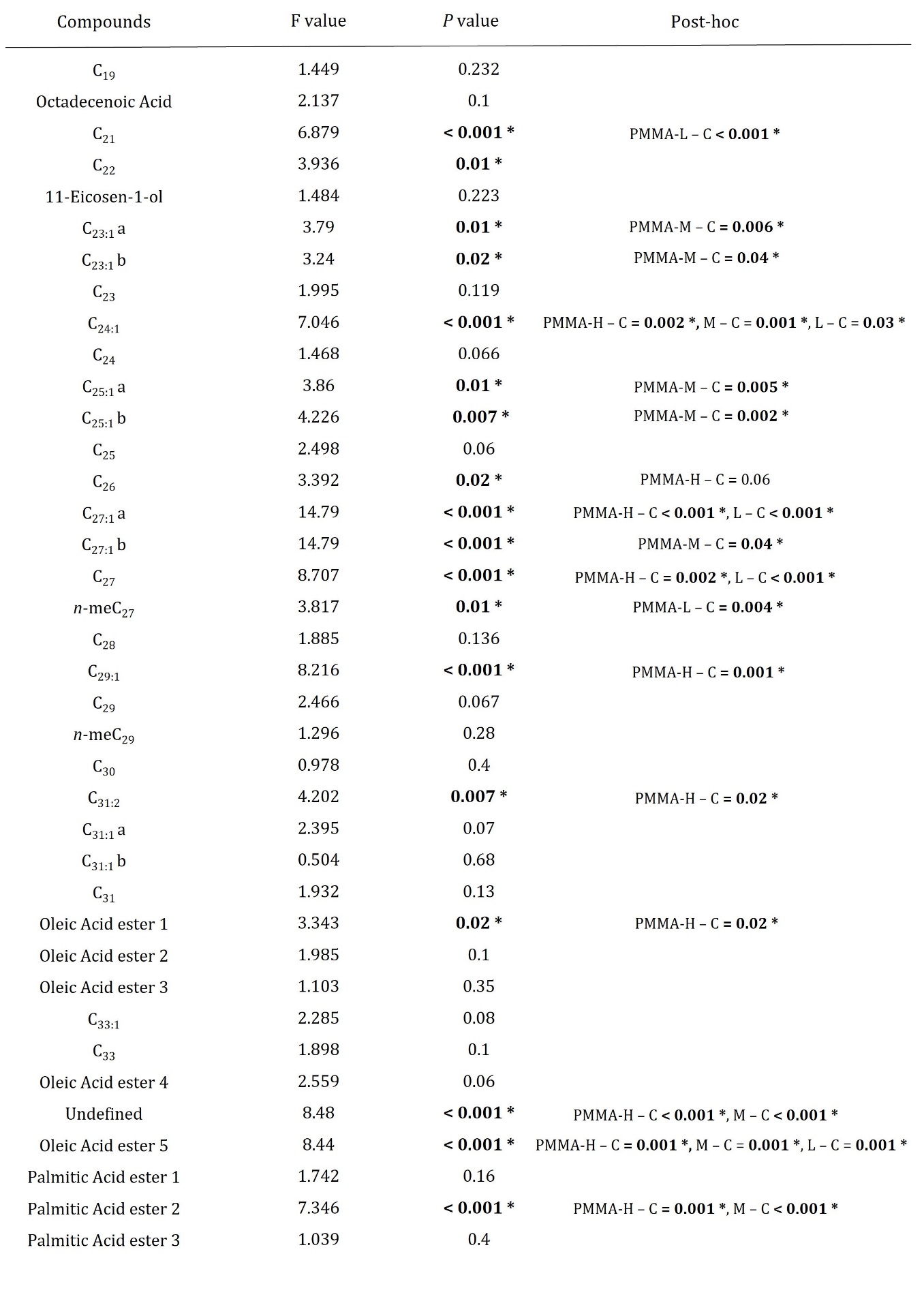

**Table SM2: Amount of cuticular hydrocarbons in bees treated with PMMA (L, M and H) compared to control bees.** Table gives F values and *p* values from ANOVA performed on each compound and *p* value from post-hoc analysis across all groups.

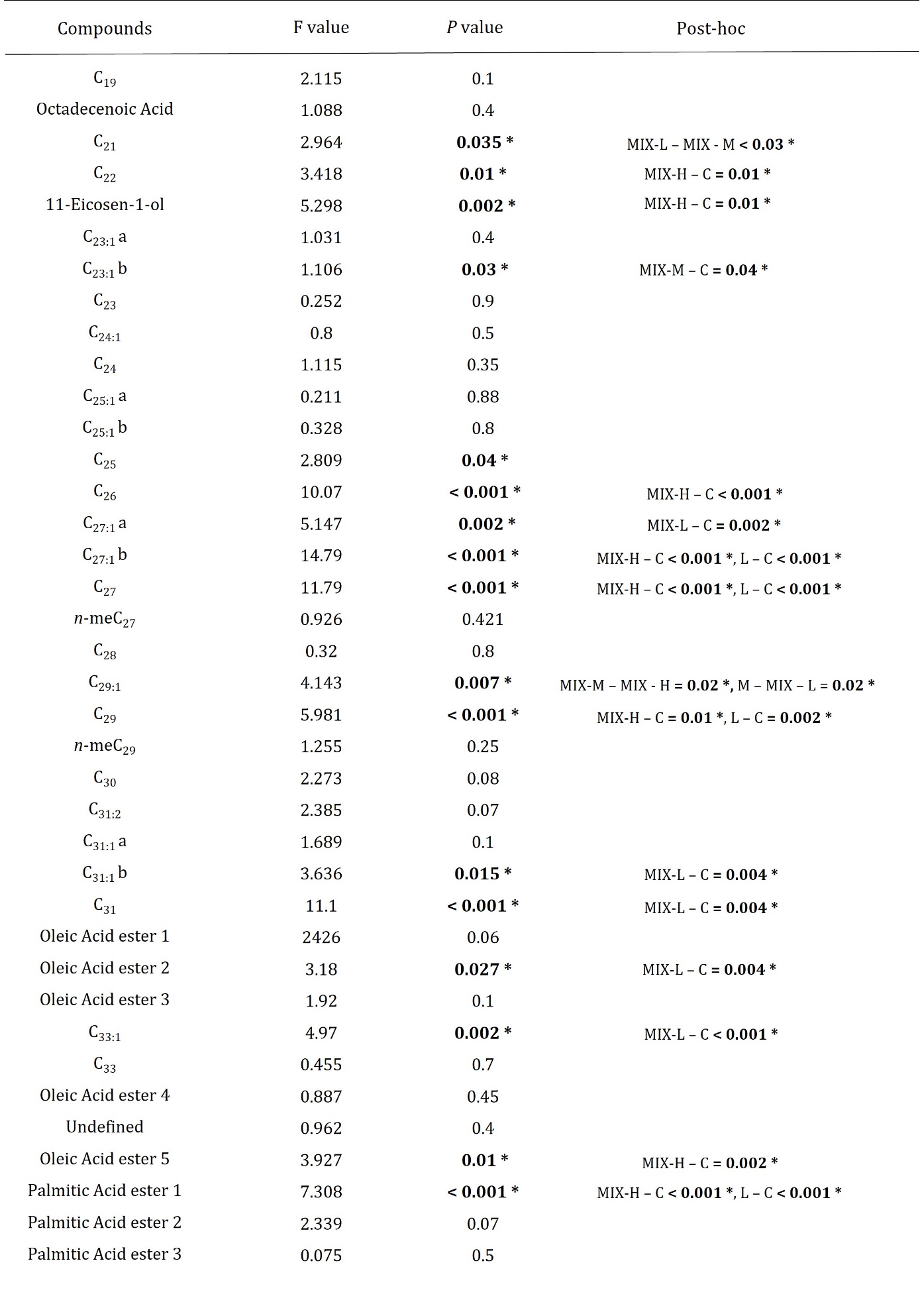

**Table SM3: Amount of cuticular hydrocarbons in bees treated with MIX (L, M and H) compared to control bees.** Table gives F values and *p* values from ANOVA performed on each compound and *p* value from post-hoc analysis across all groups.

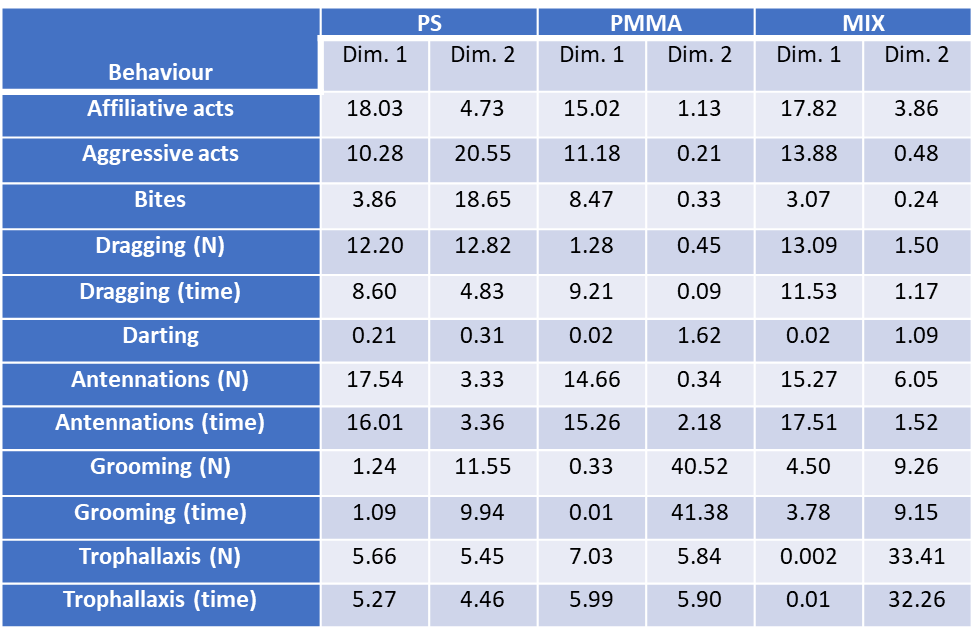

**Table SM4. Contribution of individual behaviours in the initial two dimensions of PCAs performed on bees exposed to PS, PMMA, MIX, and control bees.** The most important variables in the PCA performed on bees exposed to PS and control bees were affiliative acts (18%) and the number and duration of antennations (17.5% and 16%, respectively) for PC1, while the most important variables in PC2 were aggressive acts (20.5%), biting events (18.6%) and dragging events (12.8%). Regarding the PCA of bees exposed to PMMA, the most important variables in PC1 were affiliative acts (15%) and the number and duration of antennations (14.6% and 15.2%, respectively), while the most important ones in PC2 were the number and duration of grooming (40.5% and 41.4%, respectively) and the number and duration of trophallaxis (5.8% and 5.9%, respectively). In the PCA performed on bees exposed to MIX, affiliative acts (17.3%) and the number and duration of antennae (15.3% and 17.5%) were the most important variables in PC1, while the number and duration of trophallaxis (33.4% and 32.3%) and those of grooming (9.2% and 9.1%) were the most important in PC2.

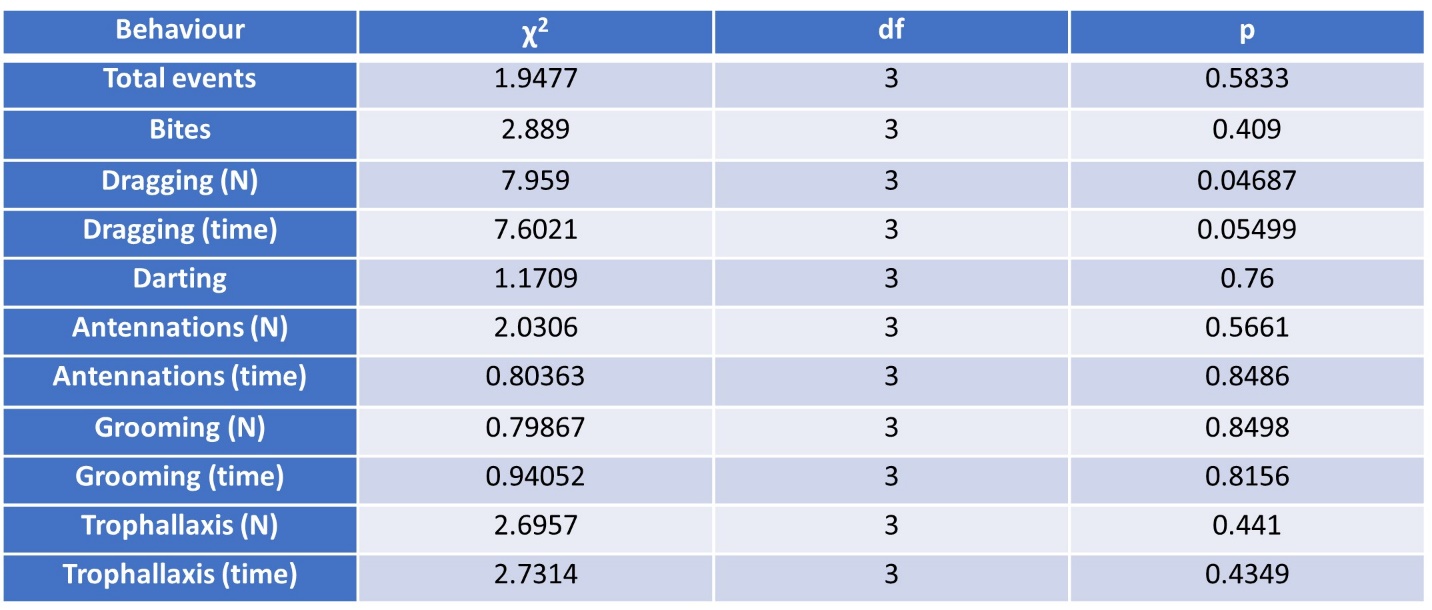

**Table SM5: Analysis of individual behaviours towards bees exposed to PS at different concentrations (PS-L, PS-M, PS-H) and control bees.** Detailed analysis of individual behaviours revealed no significant differences between groups (*p* > 0.05).

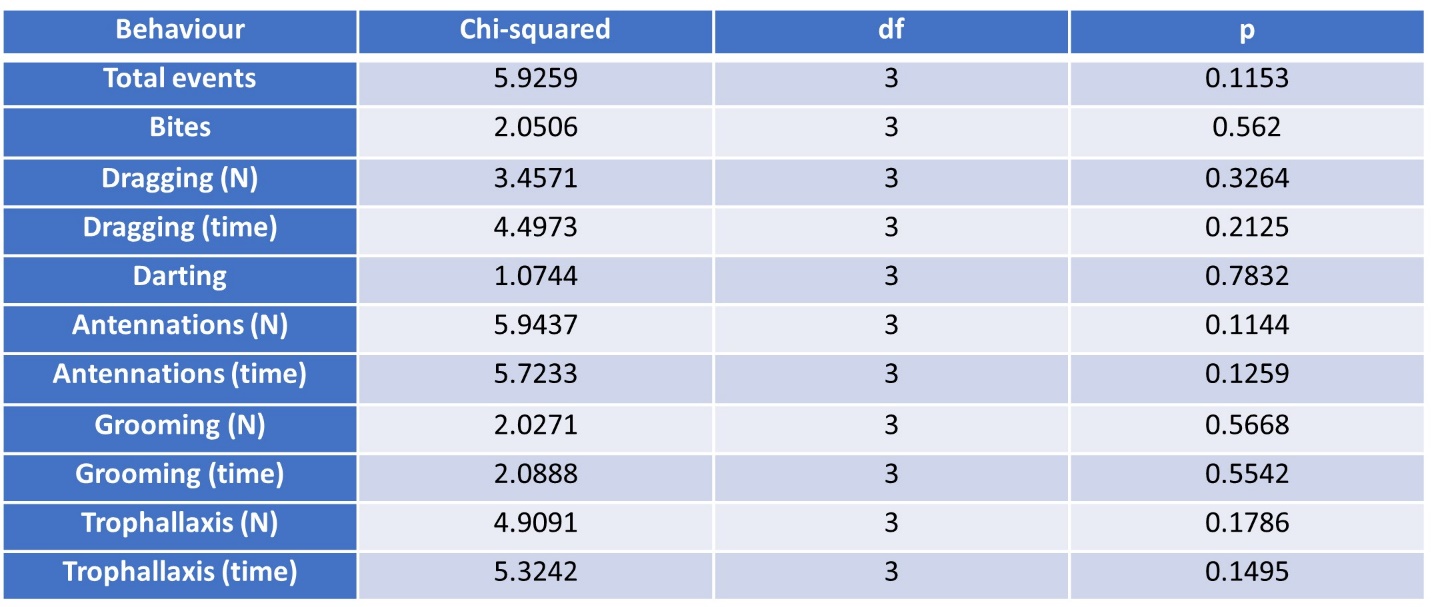

**Table SM6: Analysis of individual behaviours towards bees exposed to PMMA at different concentrations (PMMA-L, PMMA-M, PMMA-H) and control bees.** Detailed analysis of individual behaviours revealed no significant differences between groups (*p* > 0.05).

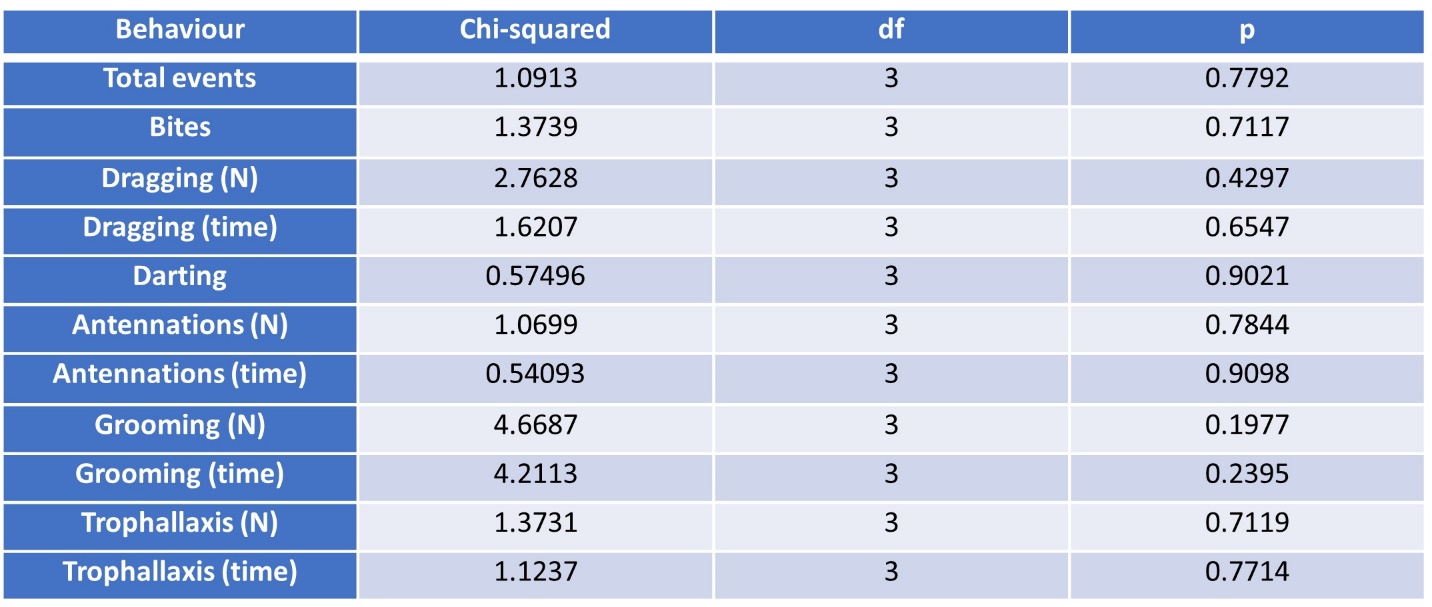

**Table SM7: Analysis of individual behaviours towards bees exposed to MIX at different concentrations (MIX-L, MIX-M, MIX-H) and control bees.** Detailed analysis of individual behaviours revealed no significant differences between groups (*p* > 0.05).
